# Rapid in situ forming PEG hydrogels for mucosal drug delivery

**DOI:** 10.1101/2024.08.16.608319

**Authors:** Taj Yeruva, Robert J. Morris, Luke Zhao, Peter Kofinas, Gregg A. Duncan

## Abstract

*In situ* gelling polymeric biomaterials have proven useful as drug delivery vehicles to enable sustained release at sites of disease or injury. However, if delivered to mucosal tissues, such as the eyes, nose, gastrointestinal, and cervicovaginal tract, these gels must also possess the ability to adhere to an epithelium coated in mucus. Towards this end, we report a new rapid *in situ* gelling polyethylene glycol-based hydrogel. Unlike other chemistries that enable rapid gel formation which form via irreversible covalent bonds, we use a bio-reducible linker allowing the gels to be naturally degraded over several days once administered. We identified a set of 6 lead formulations, which rapidly form into disulfide-linked PEG hydrogels in 30 seconds or less. These rapidly forming PEG hydrogels were also able to conform and adhere to mucosal tissues via PEG-mucin entanglements and hydrogen bonding. Controlled release of protein-based cargoes from the PEG gels was achieved over several hours whereas 40 nm nanoparticle-based cargos were retained over 24 hours. We also found these rapid in situ forming PEG gels were well-tolerated by mammalian cells. These studies support further testing and development of rapid *in situ* forming PEG gels for drug delivery to improve therapeutic retention and efficacy at mucosal sites.

## Introduction

Mucosal tissues such as the eye, stomach, nose, lung, and vagina are often sites of inflammatory and infectious diseases making them a desirable site for therapeutic delivery. Moreover, the mucosa may also be useful as a non-invasive route for drug delivery to local and distal sites as these tissues possess a large surface area that is highly vascularized and immunologically active [1–3]. However, mucosal tissues possess natural defense mechanisms to facilitate the rapid clearance of potentially infectious or irritating materials from these sites which can lead to reduced drug bioavailability when locally administered [4–7]. This has motivated use of formulations with enhanced viscosity (e.g., ointments, creams, gels) to act as a local drug depot for sustained delivery [8–10]. Thus, biomaterial systems have been sought to provide these benefits for delivery of existing drugs and others under clinical development [11].

Delivery of therapeutic-loaded gels to the cervicovaginal, ocular, nasal, and rectal routes have often necessitated design of *in situ* forming hydrogels which can be administered in a liquid form and rapidly transform upon administration to act as a local drug depot [7,12–14]. This can be achieved with thermosensitive polymers such as Pluronic and/use of click chemistry including dibenzocyclooctyne (DBCO)-azide and norbornene-tetrazine for rapid hydrogel assembly [15– 19]. A drawback of the latter approach using click chemistry is these gels are not reversible and contain crosslinks which will remain intact such that these gels will not be naturally degraded once they have delivered a therapeutic payload. Sulfhydryl crosslinked hydrogel biomaterials can also be constructed using maleimide or vinyl sulfone as reactive groups [20–22]. While highly stable, sulfhydryl crosslinked gels typically form on the order of minutes which is less than ideal for in situ gelation. As an alternative, sulfide-dipyridyl disulfide reactions can also occur rapidly but, in general, they are underexplored for hydrogel formulations used in drug delivery. These crosslinks can also be reversed under reducing conditions and are likely to degrade over time once administered due to biological tissues. Moreover, this cross-linking mirrors that found with mucus gels and prior work has shown hydrogel attachment to mucosal tissues may be facilitated via reactions with available cysteines [23–25].

In this work, we designed hydrogel biomaterials which form rapidly through thiol-pyridyl disulfide exchange. This material is composed of polyethylene glycol (PEG), a widely used polymer, and forms nearly instantaneously upon mixing under physiological conditions. Since the prepared hydrogels have thiol and pyridyl sulfides, we hypothesized adhesion of the PEG hydrogel to mucosal tissues may be favored through direct reaction with thiol groups on mucin biopolymers. To determine if potentially useful for drug delivery applications, we assessed the release rate of different types of cargo encapsulated within the PEG gels and evaluated their biocompatibility. This study establishes a new simple approach to formulate rapid in situ forming PEG gels which could be used in mucosal drug delivery applications.

## Materials and Methods

### Materials

4-arm PEG-SH (PEG-4SH) 10kD and 20kD was purchased from Laysan Bio. 4-arm PEG-OPSS (PEG-4OPSS) 10kD and 20kD was purchased from Creative PEG works. Dithiothreitol (DTT) and L-Glutathione reduced was obtained from Sigma Aldrich. Tetramethylrhodamine labeled BSA (TRITC-BSA) and Cyanine 5 conjugated IgG (Cy5-IgG) were purchased from Protein Mods. Resazurin reagent and Calcein AM were purchased from Biotium. Propidium Idodide (Invitrogen) FluoSpheres and cell culture reagents such as DMEM and fetal bovine serum were obtained from Thermo Fisher Scientific.

### Hydrogel Synthesis

4-arm thiol terminated PEG (PEG-4SH) dissolved in phosphate buffered saline (PBS) (>0.5% w/v) is mixed with equal volumes of 4-arm orthopyridyl disulfide terminated polyethylene glycol (PEG-4OPSS) dissolved in PBS (>0.5% w/v) for crosslinking and rapid gelation. Tube inversion method was used to measure gelation time at body temperature of 37 □. Briefly, polymer solutions of 100 µL each are mixed in Eppendorf tubes placed on heat block at 37□ . Tube was taken out and inverted to see gelation of the polymers every 5 sec. Time was noted when the solution stopped running on the walls of the tubes. It was repeated for all formulations of different polymer weight percentages and performed in triplicates.

### Equilibrium swelling

100 µL of cylindrical gels were made by mixing the polymer solutions in equal volumes in tip cutoff 1mL syringes. Gels were then pushed out of the syringes into 5 mL of PBS and incubated at 37□ for 24 hours to reach equilibrium swelling. After 24 hours, gels were taken out of PBS and weighed to measure mass of swollen network. Gels were then freeze dried to measure mass of dried network. Equilibrium swelling ratio was calculated using below equation.

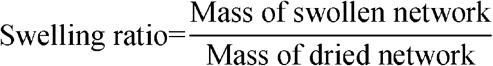

### Degradation

Degradation of PEG hydrogels were measured in PBS, PBS+ dithiothreitol (DTT) and PBS+ glutathione. To measure degradation in PBS, 100 µL cylindrical gels were prepared and placed in 5mL PBS at 37□ as described previously. Gels were weighed before immersing in PBS and at different time points after immersion in PBS. Percentage mass remaining was calculated using below equation. To measure degradation in reducing agents, 200 µL of 100 mM DTT or 100 mM glutathione in PBS was added to 100 µL gels prepared in Eppendorf tubes and incubated at

37□ . Tube was inverted every minute for DTT and every ∼ 10 hours for glutathione to check gel degradation. Time was noted when the entire gel transitioned to liquid state.

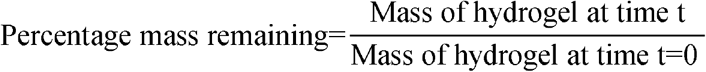

### Self-healing

100 µL square gels were formed in 3D printed square molds. Two gels were pushed out of the molds and were brought together for 10 min before holding vertically to observe self-healing. One of the gels was kept transparent while rhodamine B was added to the other gel to aid in visualization.

### Bulk rheology

To confirm successful formation of PEG hydrogels and to understand polymer weight percentages on mechanical properties, bulk rheological measurements were performed using ARES G2 rheometer (TA instruments). PEG gel precursor solutions were loaded on to 25 mm diameter parallel plate at a gap of 1000 μm at 37^º^C. Solution was allowed to gel and equilibrate to 37^º^C for 5 min. Humidity chamber was used prevent solvent evaporation and consequent hydrogel drying. To determine the linear viscoelastic region of the fully formed gel, a strain sweep measurement was performed at 0.1-10% strain at a frequency of 1 rad/s. To determine the elastic modulus, G’(ω), and viscous modulus, G’’(ω), a frequency sweep measurement is conducted within the linear viscoelastic region of the gel, at 10% strain amplitude and angular frequencies from 0.1 to 100 radians/s. To elucidate interaction between PEG solution with mucins, flow sweep was performed from a shear rate of 0.01 to 100 s^-1^ using 40 nm cone-plate geometry.

### Particle tracking microrheology

Particle tracking microrheology was performed as described in our previous work [26]. Briefly, solutions are prepared with 1 μL of ∼0.002% w/v suspension of 100 nm fluorescent muco-inert nanoparticles and added into a 25 µL solution of PEG-4SH and PEG-4OPSS prior to gelation in a custom microscopy chamber, sealed with a cover slip, and equilibrated for 30 min at room temperature before imaging. Ten-second movies at 30 ms temporal resolution were acquired with a high-speed CMOS camera equipped on an inverted confocal microscope with a 63x/1.4 NA oil objective. Movies were analyzed using custom written tracking software in MATLAB to extract 2D x, y-coordinates of MIP centroids over time. From these trajectories, time-averaged mean squared displacement (MSD; ⟨Δ*r*^2^(τ)⟩) as a function of lag time, τ, is calculated as ⟨ Δ*r*^2^(τ)⟩ = ⟨[x(t+τ)-x(t)]^2^ + ⟨[y(t+τ)-y(t)]^2^⟩ where x(t) and y(t) are the spatial coordinates of particles as a function of time *t*. Using the generalized Stokes-Einstein relation, measured MSD values are used to compute viscoelastic properties of the hydrogels (*e*.*g*., *G*’(ω), *G*’’(ω), micro-viscosity). Hydrogel network pore size, ξ, was estimated based on MSD using the equation, ξ≈(⟨ Δ*r*^2^(τ) ⟩)^1/2^ + a [27]. All measurements described were performed in n=3 PEG gel preparations per formulation.

### Drug release

0.2% w/v TRITC labeled bovine serum albumin (TRITC-BSA) or Cy5 labeled IgG (Cy5-IgG) was added to 2% w/v 4-arm PEG-SH solution and mixed with equal volumes of 3% w/v of 4-arm PEG-OPSS in tip cut-off syringes to make 100 μL cylindrical hydrogels. Resulting gels comprised of 0.1% w/v TRITC-BSA/ Cy5-IgG, 1% w/v 4-arm PEG-SH and 1.5% w/v 4-arm PEG-OPSS were immersed in 1 mL of PBS and incubated at 37□ . For nanoparticle loading, 2% v/v FluoSpheres were encapsulated. Supernatants were collected at each time point and replaced with fresh buffer. To examine protein release, fluorescence of the collected supernatants was measured by UV/Vis spectroscopy using Tecan spark multimode microplate reader. Fluorescence Ex/Em wavelengths of 545/575 was used for TRITC-BSA quantification and Ex/Em wavelengths of 650/670 was used for Cy5-IgG quantification.

### Mucoadhesion

Mucoadhesive strength of the gels was measured using a pull-apart adhesion test[28]. Square sections of 10 mm fresh porcine intestine were cut and 100 μL of PEG gel precursor solutions were applied on luminal side of the tissue. It was allowed to set for 5 min and apical side of the intestinal tissue was superglued to the clamps of a dynamic mechanical analyzer (TA Instruments, DMA Q800). Samples were initially isothermally compressed at a force of 1N for 5 minutes to ensure the superglue dries and pulled at a rate of 0.5 N/min until failure. The adhesion strength of each sample was recorded and replicated three times. To visualize mucoadhesion, a solution of 1% w/v PEG-4SH, 2% w/v PEG-4OPSS, and 0.01% w/v rhodamine B (200 µL total) was added onto a small section of intestinal tissue obtained from Animal Biotech Industries and allowed to set for 2 min before holding vertically to visualize and take photographs.

### Biocompatibility

Biocompatibility of PEG polymers and PEG gels was evaluated on HEK 293T cells. Briefly, HEK 293T cells were cultured in DMEM supplemented with 10%FBS and 1% penicillin-streptomycin. Cells were seeded onto 96 well plate at a density of 20,000 cells per well. After allowing cell adherence overnight, cells were treated with 20 μL solution of polymers dissolved in PBS pH 7.4 in the concentration range of 1% -4% w/v or 20 μL of PEG-4SH and PEG-4OPSS gels and incubated at 37□, 5% CO_2_ for 24 hours. Following the treatment, cell culture supernatant containing polymers or gels was removed and replaced with resazurin containing media. Cells with the resazurin reagent were incubated at 37, 5% CO_2_ for 3 hours. 100 μL of cell culture supernatant containing resazurin was transferred to a 96-well black plate and fluorescence was measured at Ex/Em wavelengths of 570/585 nm. Cell viability was measured relative to the cells grown in media without any treatment. For live/dead staining, í 10^5^ cells were seeded in an 8-well chambered cell culture slide. After allowing cell adherence overnight, 50 μL of 2% PEG-4SH and 2% PEG-4OPSS gels were added to each well and incubated at 37□, 5% CO_2_ for 24 hours. Following the treatment, live cell staining was performed using 1μ? calcein AM and 500 nM propidium iodide and cells were imaged using Zeiss confocal microscope.

## Results and Discussion

### Formulation of rapid in-situ forming PEG hydrogels

In this study, we developed a new method for rapid formation of polyethylene glycol (PEG) based hydrogels at physiological conditions of 37°C and pH of 7.4. These gels are also formed via bio-reducible disulfide linkers by combining thiol terminated 4-arm PEG (PEG-SH) and OPSS terminated 4-arm PEG (PEG-OPSS). An overview of crosslinking mechanism for 4-arm PEG-SH and 4-arm PEG-OPSS is depicted in **Figure 1**. After being dissolved in phosphate buffered solution (PBS), we observed gel formation within 30 seconds through rapid crosslinking via di-sulfide bond formation. **Table 1** summarizes the time to gel as well as degradation time under reducing conditions for 6 lead formulations. Although a wide range of weight percentages were tested and found to rapidly form gels as noted in Table S1, lead formulations were selected which retained stability in PBS for ≥ 24 hours.

**Table 1.** Gelation and degradation times of rapid forming PEG gels.

| Formulation number | 4-arm PEG-SH (% w/v) | 4-arm PEG-OPSS (% w/v) | Gelation time | Degradation time in DTT | Degradation time in GSH |
| --- | --- | --- | --- | --- | --- |
| 10kD PEG |  |  |  |  |  |
| F1 | 1 | 1.5 | ~15 s | ~ 8 min | ~ 30 hr. |
| F3 | 1 | 2 | ~ 5 s | ~ 9 min | ~ 60 hr. |
| F5 | 2 | 2 | ~ 5 s | ~ 6 min | ~ 60 hr. |
| 20kD PEG |  |  |  |  |  |
| F2 | 1 | 1.5 | ~ 30 s | ~ 3 min | ~ 30 hr. |
| F4 | 1 | 2 | ~ 20 s | ~ 5 min | ~ 30 hr. |
| F6 | 2 | 2 | ~ 15 s | ~ 7 min | ~ 40 hr. |

**Figure 1.**
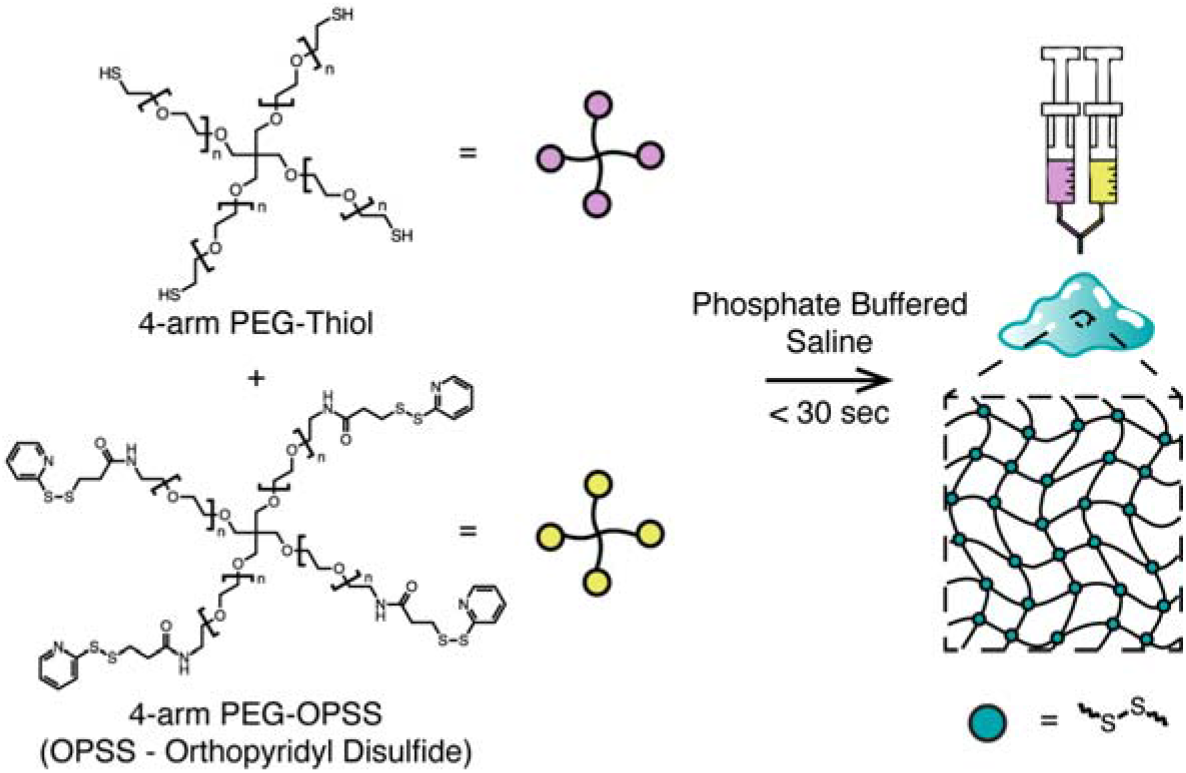
Schematic illustrating the preparation of rapid in situ forming PEG hydrogels.

The disulfide reducing agent glutathione (GSH) is present throughout tissues in the body at a µM to mM concentration range [29]. Thus, the degradation rate of PEG gels once administered *in vivo* is likely to be affected by the presence of GSH. Further, external disulfide reducing agents such as dithiothreitol (DTT) can be used for removal of the gel from the site of application as needed. We tested the effect of these compounds on our PEG gels *in vitro* using 100 mM DTT and GSH individually. As indicated in **Table 1**, DTT degrades gels under 10 minutes whereas glutathione degrades in 1 to 3 days.

### Characterization of rapid forming PEG gels

Effect of polymer weight percentage and polymer molecular weight on mechanical strength of hydrogels are studied in **Fig. 2A**. Formulations F1 to F5 have a storage modulus ranging from ∼400–800 Pa. Formulation 6 containing 20 kDa sized 2% w/v 4-arm PEG-SH and 2% w/v 4-arm PEG-OPSS possessed a significantly higher storage modulus of 1712 ± 267 Pa, indicative of an increased elasticity in comparison to other formulations tested.

**Figure 2.**
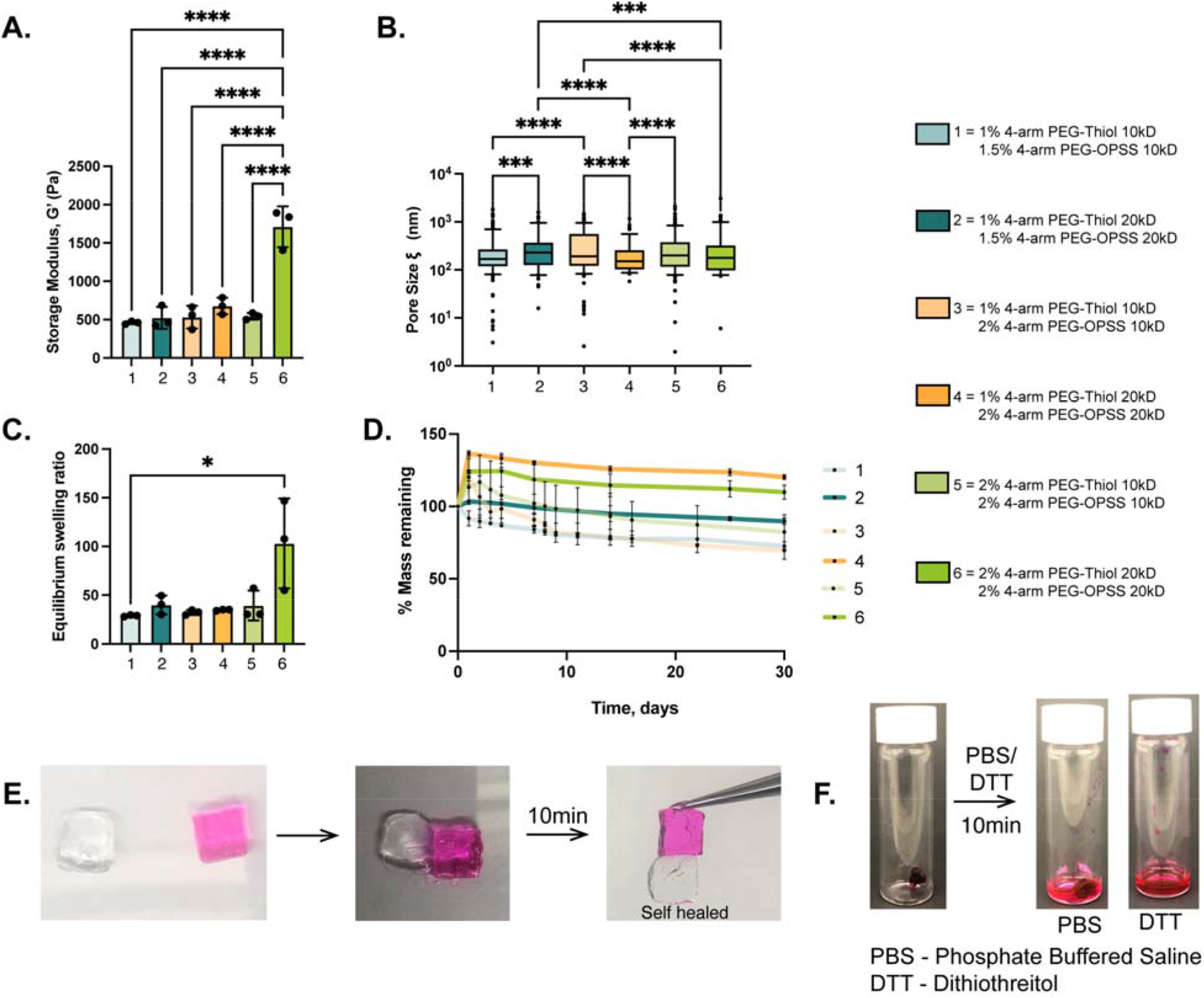
Physical characterization of rapid in situ forming PEG gels. (**A**) Storage modulus (G’) for each gel at frequency of 1 Hz. (n=3), \*\*\*\**p* <0.0001 for one-way ANOVA with Tukey’s multiple comparison test. (**B**) Estimated pore size size for particle tracking microrheology using densely PEGylated 100 nm nanoparticles as probes. * *p* <0.05, \*\*\**p* <0.001, \*\*\*\**p* <0.0001 for Kruskal-Wallis test. (**C**) Equilibrium swelling ratio after immersion in PBS. (n=3), \**p* <0.05 for Kruskal-Wallis test (**D**) Degradation profile of PEG gels after immersion in PBS for 30 hours. (**E**) Photographs of self-healing behavior for PEG gels. (**F**) Dissolution of PEG gels after 10 minutes in PBS or under reducing conditions in DTT. Data shows mean□±□SD

We next evaluated the pore size of each formulation using particle tracking microrheology (**Fig. 2B)**. We hypothesized increasing PEG-SH/PEG-OPSS concentrations or molecular weights would lead to decreases in pore size which may change release kinetics of encapsulated cargoes. Interestingly, we found only a weak dependence on total PEG concentration with a relatively small range of pore sizes observed across formulation conditions. We did observe changes in pore size as a function of PEG MW, but a consistent trend was not observed. Total available functional groups for crosslinking, flexibility of polymer chain to facilitate disulfide bond formation and steric hindrance due to bulkiness of OPSS reactive groups compared to SH could be reasons for inconsistent trends.

Swelling and degradation dictate the rate of release of therapeutics and biodegradability, which are vital for use in biomedical applications. **Fig. 2C** shows the equilibrium swelling ratio for each formulation. F6 with highest storage modulus also has the highest swelling ratio and is 2-fold higher than rest of the formulations. For degradation studies (**Fig. 2D**), PEG gels are immersed in PBS volume that is 50 times the volume of gel. An initial increase in gel mass was observed due to swelling followed by slow degradation. Gels made with 10kD polymer, F1, F3, and F5 showed degradation up to 30% of initial swollen mass in 30 days whereas gels made of 20kD polymers, F2, F4, and F6 degraded up to 15% of initial swollen mass in the same time frame. Longer retention and slow degradation of PEG gels can be due to stability of ether linkages of PEG polymer and susceptibility to degradation due to auto-oxidation [30].

**Fig. 2E** illustrates the self-healing nature of the rapid forming PEG gels. When two gels of formulation F2, 1% w/v 4-arm PEG-SH, 20kD crosslinked with 1.5% w/v 4-arm PEG-OPSS, 20kD are brought together, they self-heal at the site of attachment through disulfide and hydrogen bond formation. Self-healing hydrogels can repair in the event of fracture and restore to original mechanical properties which helps in prevention of detachment at the site of application and leakage of therapeutic cargo. **Figure 2F** further illustrates the linkers within the PEG gels are reducible as immersion of the rhodamine-loaded PEG gel in PBS alone shows rhodamine release but no degradation of the gel. However, addition of DTT shows complete degradation of the gel within 10 minutes.

### Mucoadhesive properties of rapid in situ forming PEG hydrogels

We next evaluated the ability of these PEG gels to adhere to mucosal tissues. Prior work has shown PEG based biomaterials can adhere to mucus-coated tissues through a combination of PEG-mucin entanglement and hydrogen bonding (**Fig. 3A**) [31–33]. We hypothesized our gels may also be able to form disulfide bonds with cysteine-rich domains of mucins via PEG-OPSS and/or PEG-SH leading to increases in mucoadhesion. To test this, we applied PEG-4OPSS and PEG-4SH solutions to the surface of pig intestine where instantaneously formed a uniform layer of PEG gel (**Fig. 3B**). Pull-apart tests were then performed to measure their mucoadhesive strength immediately and 24 hours after application (**Fig. 3C and 3D**). We also included for comparison tissues treated with 2% w/v 4-arm PEG-DBCO and 2 % w/v 4-arm PEG-azide as a control for PEG gels without OPSS/SH groups to form disulfide bonds and 4% w/v chitosan solution which is known to possess mucoadhesive properties. All formulations tested were found to adhere to pig intestine with mucoadhesive strengths >600 Pa. Although statistically non-significant, the 4% w/v chitosan possessed greater mucoadhesive strength than all PEG gel formulations likely due to the net-positively charged chitosan adhering to net-negatively charged mucin chains. However, PEG-DBCO and PEG-azide gels showed comparable mucoadhesive strength to PEG-OPSS and PEG-SH gels indicating our gels are not likely able to access the cysteines on the mucosal tissue. Therefore, adhesion is likely mediated by entanglement during sol-gel transition and hydrogen bonding. This is further confirmed through flow sweep measurements, which showed no change in viscosity when mucin solutions were mixed PEG-4SH and PEG-4OPSS indicating no covalent bond formation **(Fig. S2C)**.

**Figure 3.**
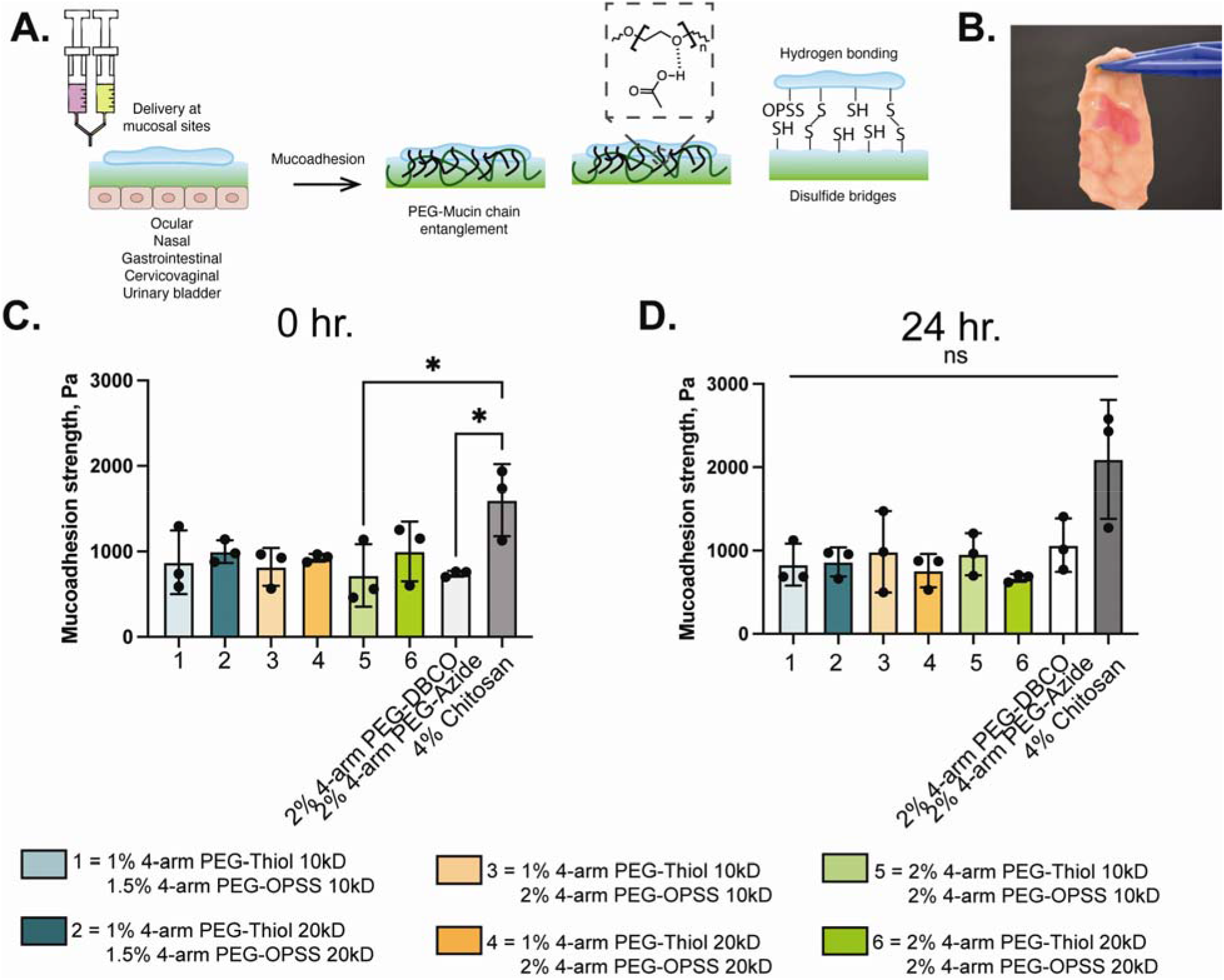
Mucoadhesive properties of rapid in situ forming PEG hydrogels. (**A**) Schematic illustrating potential mechanisms enabling PEG gel adhesion to mucosal tissues. (**B**) Image of PEG gels following application to porcine intestinal tissue. (**C**) Mucoadhesive strength as measured by pull-apart test for PEG gels & 4% w/v chitosan immediately (n=3), * *p* <0.05 for on-way ANOVA with Tukey’s multiple comparison test and (D) 24 hours (n=3), non-significant (ns) as per Kruskal-Wallis test after application to porcine intestinal tissue. Data shows mean□±□SD.

### Release of cargo from rapid forming PEG gels

We characterized the release kinetics of different model cargos, including bovine serum albumin (BSA), immunoglobulin (IgG), and 40 nm nanoparticles (NP), from PEG gels over 24 hours. We found BSA and IgG were released in roughly 4 hours and 10 hours respectively whereas 40 nm NP were retained within the gel for at least 24 hours **(Fig. 4)**. This is likely explained with the difference in cargo size with BSA and IgG being ∼4-5 times smaller than 40 nm NP. These data indicate release of protein therapeutics would likely be driven by diffusion out of the gel whereas delivery of encapsulated nanoparticles is more likely to be driven by gel degradation. Further, effect of polymer weight percentage and swelling ratio has no effect on BSA release, minimal effect on IgG release and highest effect on 20 nm nanoparticles exhibiting increased release in formulation 6 due to higher swelling ratio **(Fig. S5)**.

**Figure 4.**
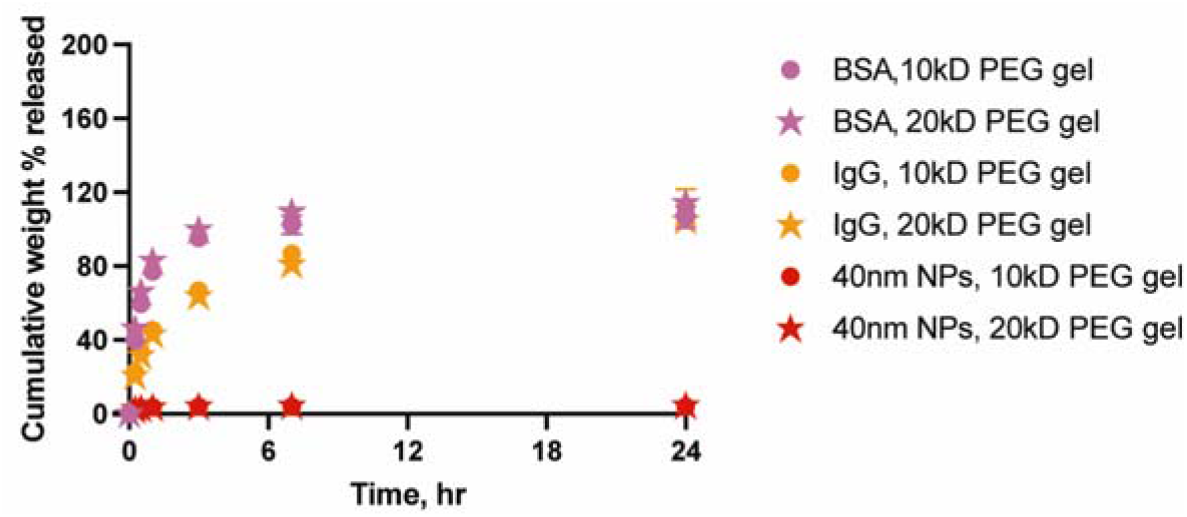
Release kinetics of model therapeutic cargo from rapid forming PEG hydrogels. Cumulative mass release profiles of bovine serum albumin (BSA), immunoglobulin (IgG), and 40 nm nanoparticles (NP) from PEG hydrogels with either 10 kDa or 20 kDA PEG-SH and PEG-OPSS over 24 hours (n=3). Data shows mean□±□SD.

### Biocompatibility of rapid forming PEG gels

To ensure the gel precursor components and the PEG gel itself are well-tolerated by mammalian cells, we conducted biocompatibility studies using HEK-293 cells (**Fig. 5**). First, we treated HEK-293 cells with either 4-arm PEG-SH or PEG-OPSS at concentrations up to 4% w/v (**Fig. 5A**). We found 10 kDa 4-arm PEG-SH was well-tolerated at all concentrations tested. Although statistically significant differences were observed with other treatments, more than 80% cell viability was observed except for 10 kDa 4-arm PEG OPSS at 4% w/v concentration. We next treated HEK-293 cells with PEG gels and no toxicity was observed for all lead formulations tested (**Fig. 5B**). These data suggest that PEG gels are generally safe for use as an injectable biomaterial. However, acute exposure to 10 kDa 4-arm PEG formulations may pose some toxicity concerns and thus, 20 kDa PEG formulation may be better suited for use in future drug delivery applications.

**Figure 5.**
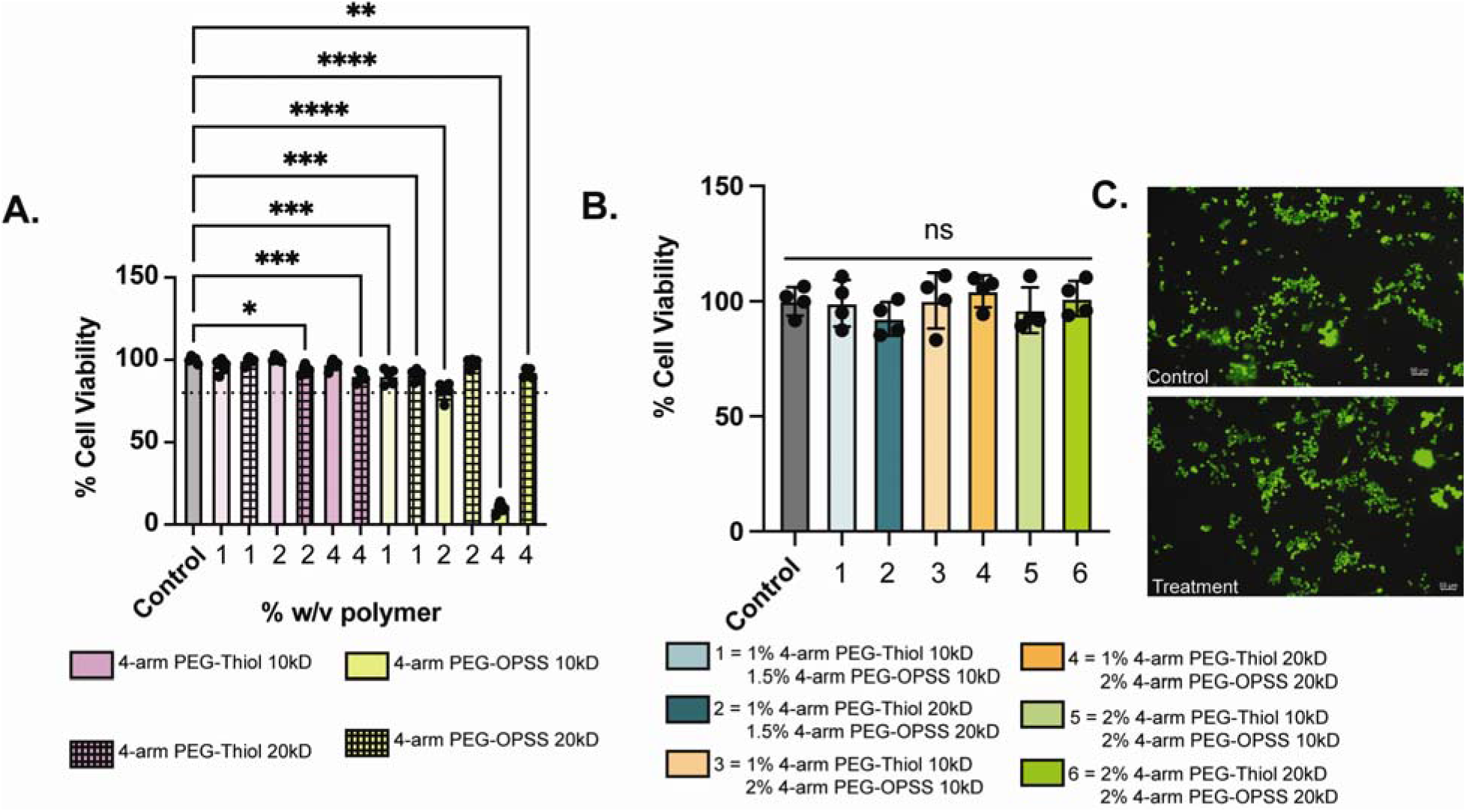
Biocompatibility of rapid forming PEG gels. Viability of HEK-293 cells following treatment with **(A)** 4-arm PEG solutions (n=5) and **(B)** PEG hydrogels (n=4) (C) Live/Dead fluorescent staining following treatment with PEG hydrogels. * *p* <0.05, \*\**p* <0.01, \*\*\**p* <0.001, and \*\*\*\**p* <0.0001 for oneway ANOVA with Dunnett’s multiple comparison test. Data shows mean□±□SD.

## Conclusion

We have developed a rapid *in situ* forming PEG hydrogel capable of adhering to mucosal tissues. Unlike other chemistries used in previous work, we can form bio-reducible disulfide linked PEG gels which are able to be degraded over time (days to weeks) upon administration. Both protein and nanoparticle-based therapeutics may be encapsulated into the gel for extended release at mucosal sites. Future work will focus on further development of PEG gels as a novel sprayable and/or injectable in situ forming hydrogel for delivery of therapeutics to mucosal tissues.

## Acknowledgements

This study was supported by NIH R21 EB030834 and NIGMS R01GM141132.

## References

[1] J.R. McGhee, J. Mestecky, M.T. Dertzbaugh, J.H. Eldridge, M. Hirasawa, H. Kiyono, The mucosal immune system: from fundamental concepts to vaccine development, Vaccine 10 (1992) 75–88. 10.1016/0264-410x(92)90021-b.

[2] E.A. Naumova, T. Dierkes, J. Sprang, W.H. Arnold, The oral mucosal surface and blood vessels, Head Face Med 9 (2013) 8. 10.1186/1746-160X-9-8.

[3] L.-A. Keller, O. Merkel, A. Popp, Intranasal drug delivery: opportunities and toxicologic challenges during drug development, Drug Delivery and Translational Research 12 (2022) 735–757. 10.1007/s13346-020-00891-5.

[4] N.A. Peppas, J.J. Sahlin, Hydrogels as mucoadhesive and bioadhesive materials: a review, Biomaterials 17 (1996) 1553–1561.

[5] T. Yeruva, S. Yang, S. Doski, G.A. Duncan, Hydrogels for Mucosal Drug Delivery, ACS Appl. Bio Mater. 6 (2023) 1684–1700. 10.1021/acsabm.3c00050.

[6] X. Murgia, B. Loretz, O. Hartwig, M. Hittinger, C.-M. Lehr, The role of mucus on drug transport and its potential to affect therapeutic outcomes, Advanced Drug Delivery Reviews 124 (2018) 82–97. 10.1016/j.addr.2017.10.009.

[7] S.R. Van Tomme, G. Storm, W.E. Hennink, In situ gelling hydrogels for pharmaceutical and biomedical applications, International Journal of Pharmaceutics 355 (2008) 1–18. 10.1016/j.ijpharm.2008.01.057.

[8] A. Bak, M. Ashford, D.J. Brayden, Local delivery of macromolecules to treat diseases associated with the colon, Adv Drug Deliv Rev 136–137 (2018) 2–27. 10.1016/j.addr.2018.10.009.

[9] I. Seah, X.J. Loh, X. Su, A topical gel for extended ocular drug release, Nature Biomedical Engineering 4 (2020) 1024–1025. 10.1038/s41551-020-00645-1.

[10] M.S. Roberts, H.S. Cheruvu, S.E. Mangion, A. Alinaghi, H.A.E. Benson, Y. Mohammed, A. Holmes, J. Van Der Hoek, M. Pastore, J.E. Grice, Topical drug delivery: History, percutaneous absorption, and product development, Advanced Drug Delivery Reviews 177 (2021) 113929. 10.1016/j.addr.2021.113929.

[11] A. Mandal, J.R. Clegg, A.C. Anselmo, S. Mitragotri, Hydrogels in the clinic, Bioengineering & Transla Med 5 (2020) e10158. 10.1002/btm2.10158.

[12] M. Agrawal, S. Saraf, S. Saraf, S.K. Dubey, A. Puri, U. Gupta, P. Kesharwani, V. Ravichandiran, P. Kumar, V.G.M. Naidu, U.S. Murty Ajazuddin, A. Alexander, Stimuli-responsive In situ gelling system for nose-to-brain drug delivery, Journal of Controlled Release 327 (2020) 235–265. 10.1016/j.jconrel.2020.07.044.

[13] M.H. Asfour, S.H. Abd El-Alim, G.E.A. Awad, A.A. Kassem, Chitosan/β-glycerophosphate in situ forming thermo-sensitive hydrogel for improved ocular delivery of moxifloxacin hydrochloride, European Journal of Pharmaceutical Sciences 167 (2021) 106041. 10.1016/j.ejps.2021.106041.

[14] Y.C. Kim, M.D. Shin, S.F. Hackett, H.T. Hsueh, R. Lima e Silva, A. Date, H. Han, B.-J. Kim, A. Xiao, Y. Kim, L. Ogunnaike, N.M. Anders, A. Hemingway, P. He, A.S. Jun, P.J. McDonnell, C. Eberhart, I. Pitha, D.J. Zack, P.A. Campochiaro, J. Hanes, L.M. Ensign, Gelling hypotonic polymer solution for extended topical drug delivery to the eye, Nature Biomedical Engineering 4 (2020) 1053–1062. 10.1038/s41551-020-00606-8.

[15] K.S. Anseth, H.-A. Klok, Click chemistry in biomaterials, nanomedicine, and drug delivery, Biomacromolecules 17 (2016) 1–3.

[16] H. Zhan, H. de Jong, D.W. Löwik, Comparison of bioorthogonally cross-linked hydrogels for in situ cell encapsulation, ACS Applied Bio Materials 2 (2019) 2862–2871.

[17] T. Yeruva, C.H. Lee, Enzyme Responsive Delivery of Anti-Retroviral Peptide via Smart Hydrogel, AAPS PharmSciTech 23 (2022) 234. 10.1208/s12249-022-02391-w.

[18] C. Bahou, R.J. Spears, A.M. Ramírez Rosales, L.N.C. Rochet, L.J. Barber, K.S. Stankevich, J.F. Miranda, T.C. Butcher, A.M. Kerrigan, V.K. Lazarov, W. Grey, V. Chudasama, C.D. Spicer, Hydrogel Cross-Linking via Thiol-Reactive Pyridazinediones, Biomacromolecules 24 (2023) 4646–4652. 10.1021/acs.biomac.3c00290.

[19] H.T. Hoang, S.-H. Jo, Q.-T. Phan, H. Park, S.-H. Park, C.-W. Oh, K.T. Lim, Dual pH-/thermo-responsive chitosan-based hydrogels prepared using “click” chemistry for colon-targeted drug delivery applications, Carbohydrate Polymers 260 (2021) 117812. 10.1016/j.carbpol.2021.117812.

[20] T.S. Hebner, B.E. Kirkpatrick, B.D. Fairbanks, C.N. Bowman, K.S. Anseth, D.S. Benoit, Radical Mediated Degradation of Thiol–Maleimide Hydrogels, Advanced Science (2024) 2402191.

[21] J.I. Paez, A. Farrukh, R. Valbuena-Mendoza, M.K. Włodarczyk-Biegun, A. Del Campo, Thiol-methylsulfone-based hydrogels for 3D cell encapsulation, ACS Applied Materials & Interfaces 12 (2020) 8062–8072.

[22] J. Yu, X. Xu, F. Yao, Z. Luo, L. Jin, B. Xie, S. Shi, H. Ma, X. Li, H. Chen, In situ covalently cross-linked PEG hydrogel for ocular drug delivery applications, International Journal of Pharmaceutics 470 (2014) 151–157. 10.1016/j.ijpharm.2014.04.053.

[23] A. Bernkop-Schnürch, Thiomers: A new generation of mucoadhesive polymers, Advanced Drug Delivery Reviews 57 (2005) 1569–1582. 10.1016/j.addr.2005.07.002.

[24] R.P. Brannigan, V.V. Khutoryanskiy, Progress and current trends in the synthesis of novel polymers with enhanced mucoadhesive properties, Macromolecular Bioscience 19 (2019) 1900194.

[25] V. Grabovac, D. Guggi, A. Bernkop-Schnürch, Comparison of the mucoadhesive properties of various polymers, Adv Drug Deliv Rev 57 (2005) 1713–1723. 10.1016/j.addr.2005.07.006.

[26] K. Joyner, D. Song, R.F. Hawkins, R.D. Silcott, G.A. Duncan, A rational approach to form disulfide linked mucin hydrogels, Soft Matter 15 (2019) 9632–9639. 10.1039/c9sm01715a.

[27] K. Joyner, S. Yang, G.A. Duncan, Microrheology for biomaterial design, APL Bioengineering 4 (2020) 041508. 10.1063/5.0013707.

[28] J.L. Daristotle, M. Erdi, L.W. Lau, S.T. Zaki, P. Srinivasan, M. Balabhadrapatruni, O.B. Ayyub, A.D. Sandler, P. Kofinas Biodegradable, Tissue Adhesive Polyester Blends for Safe, Complete Wound Healing, ACS Biomater. Sci. Eng. 7 (2021) 3908–3916. 10.1021/acsbiomaterials.1c00865.

[29] G. Wu, J.R. Lupton, N.D. Turner, Y.-Z. Fang, S. Yang, Glutathione Metabolism and Its Implications for Health, The Journal of Nutrition 134 (2004) 489–492. 10.1093/jn/134.3.489.

[30] M.E. Payne, O.O. Kareem, K. Williams-Pavlantos, C. Wesdemiotis, S.M. Grayson, Mass spectrometry investigation into the oxidative degradation of poly (ethylene glycol), Polymer Degradation and Stability 183 (2021) 109388.

[31] L. Serra, J. Doménech, N.A. Peppas, Design of poly(ethylene glycol)-tethered copolymers as novel mucoadhesive drug delivery systems, European Journal of Pharmaceutics and Biopharmaceutics 63 (2006) 11–18. 10.1016/j.ejpb.2005.10.011.

[32] N.V. Efremova, Y. Huang, N.A. Peppas, D.E. Leckband, Direct Measurement of Interactions between Tethered Poly(ethylene glycol) Chains and Adsorbed Mucin Layers, Langmuir 18 (2002) 836–845. 10.1021/la011303p.

[33] P. Schattling, E. Taipaleenmäki, Y. Zhang, B. Städler, A polymer chemistry point of view on mucoadhesion and mucopenetration, Macromolecular Bioscience 17 (2017) 1700060.

