## Supporting Information for "Rapid in situ forming PEG hydrogels for mucosal drug delivery"

*Table S1.* Gelation and degradation times of rapid forming PEG gels.

| 4-arm PEG-SH (%w/v) | 4-arm PEG-SH (%w/v) | Gelation time | |
|---|---|---|---|
| | | 10kD PEG | 20 kD PEG |
| 0.5 | 0.5 | ~4 min | ~1.5 min |
| 0.5 | 1 | ~1.5 min | ~50 s |
| 1 | 0.5 | ~2 days | ~5 min |
| 1 | 1 | ~30 s | ~30 s |
| 1.5 | 1 | ~30 s | ~25 s |
| 1.5 | 1.5 | ~10 s | ~15 s |
| 2 | 1 | >2 hr. | ~20s |

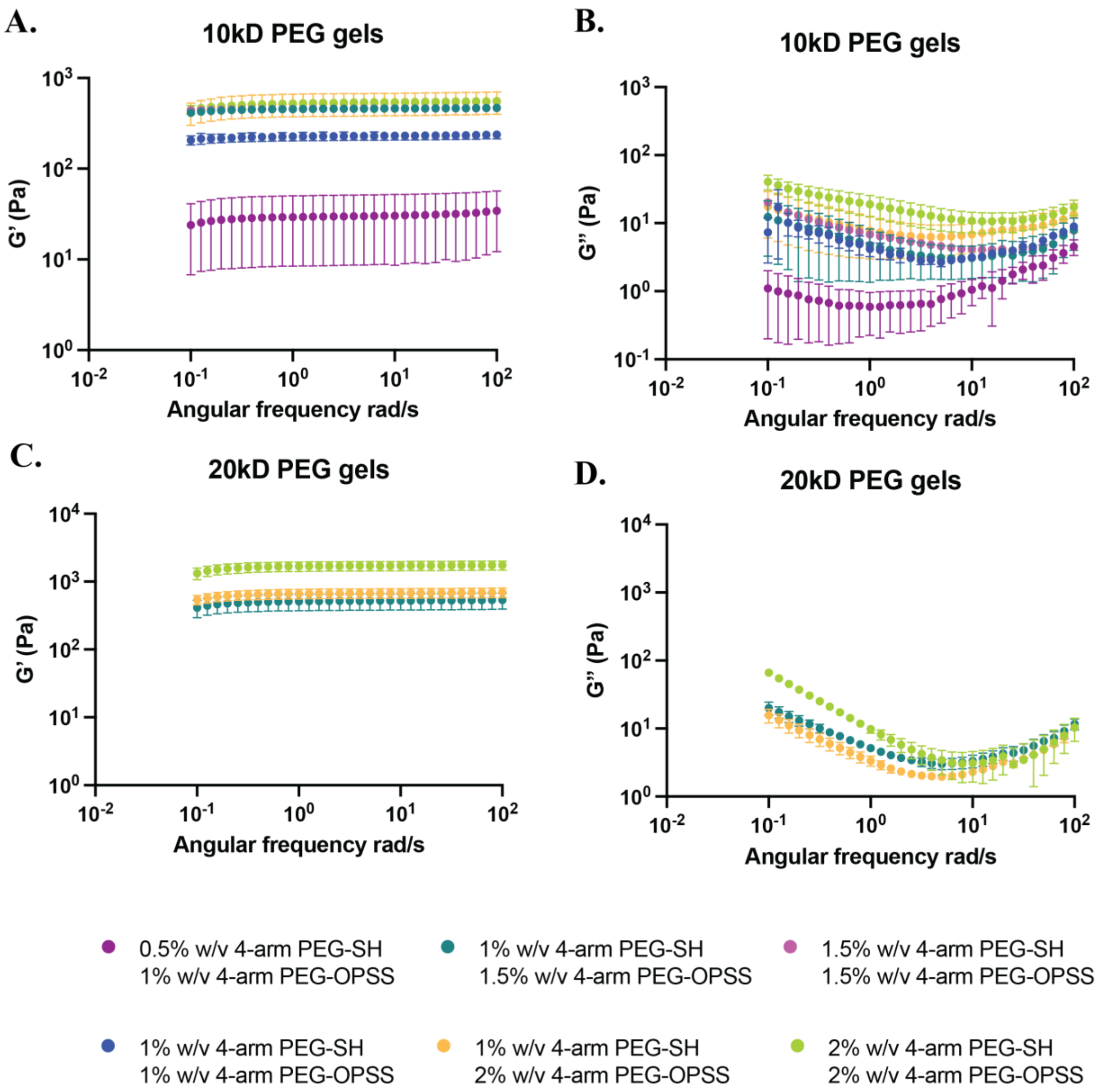


Figure S1. Bulk rheological properties of rapid in situ forming PEG gels. (A), (B) Storage modulus (G') and (C), (D) Loss modulus (G") as a function of frequency at 10% strain (n=3).

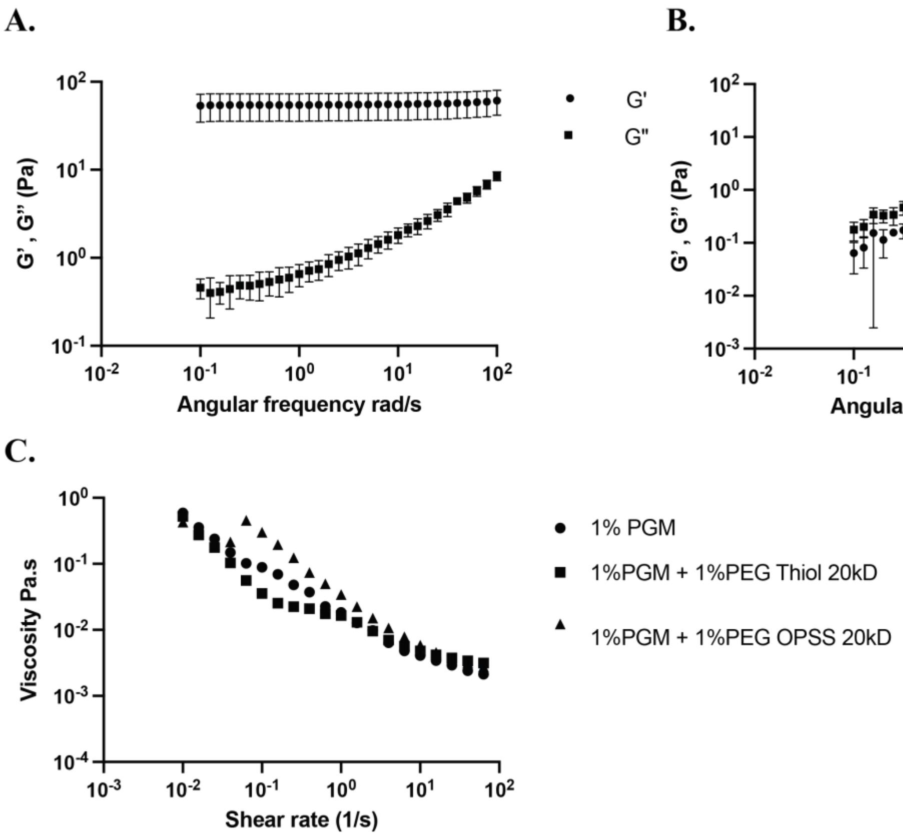


Figure S2. Bulk rheological properties of control gels used for mucoadhesion. Storage modulus (G') and Loss modulus (G") as a function of frequency at 10% strain of (A) 2% w/v 4-arm PEG-DBCO 10kD crosslinked with 2%w/v 4-arm PEG-Azide 10kD (n=3), (B) 4% w/v chitosan (n=3). (C) Flow sweep (n=1) showing no instant crosslinking between mucin chains and PEG-Thiol or PEG-OPSS indicating mucoadhesion at 0 hr. is driven by polymer-mucin chain entanglement and hydrogen bonding.

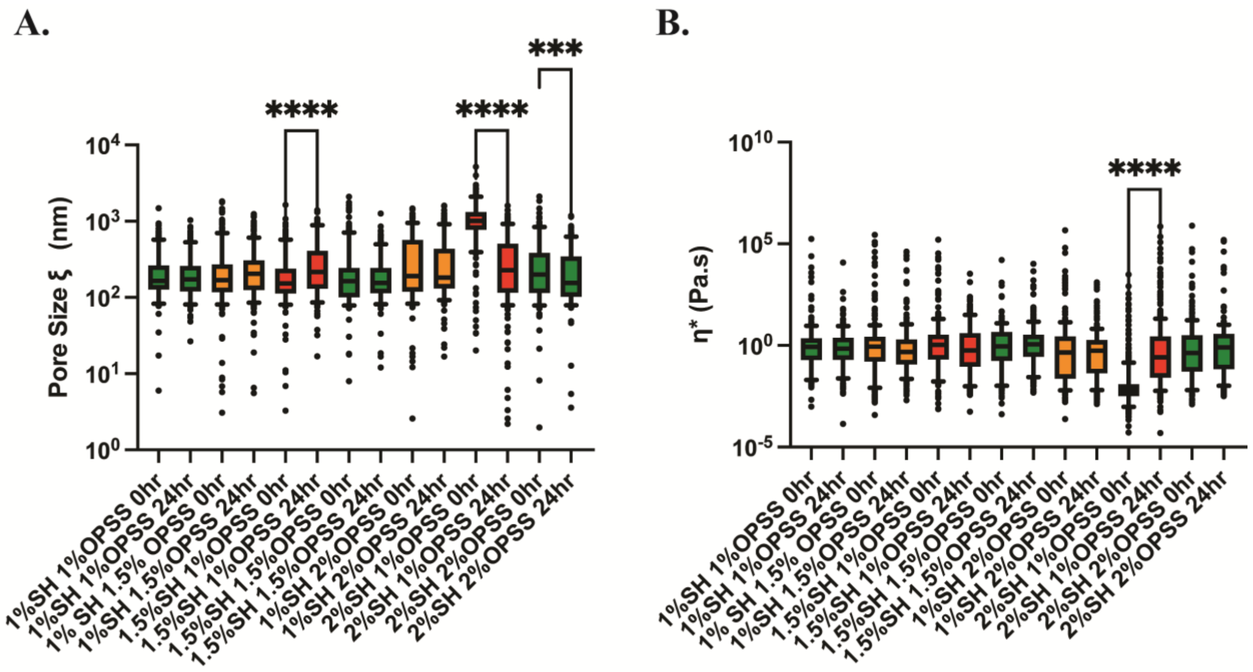


Figure S3. Micro rheological properties of 10kD PEG gels at 0 hr. and 24 hr. after mixing PEG-SH and PEG-OPSS solutions. (A) Estimated pore size based on analysis of mean square displacement (MSD) at $\tau$=1s. (B) Complex microviscosity ($\eta^*$) at a frequency $\omega = 1$ Hz calculated from measured MSD. * $p$ <0.05, ** $p$ <0.01, *** $p$ <0.001, **** $p$ <0.0001 for Kruskal-Wallis test.

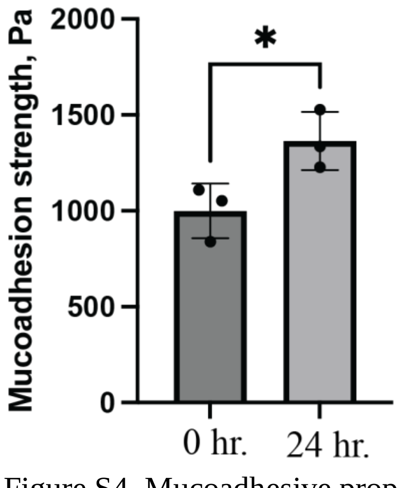


Figure S4. Mucoadhesive properties of 0.5% w/v 4-arm PEG-SH 10kD and 1% w/v 4-arm PEG-OPSS 10kD gels at 0 hr. and 24hr. after application to porcine intestinal tissue (n=3). * $p$ <0.05 for Welch’s t test.

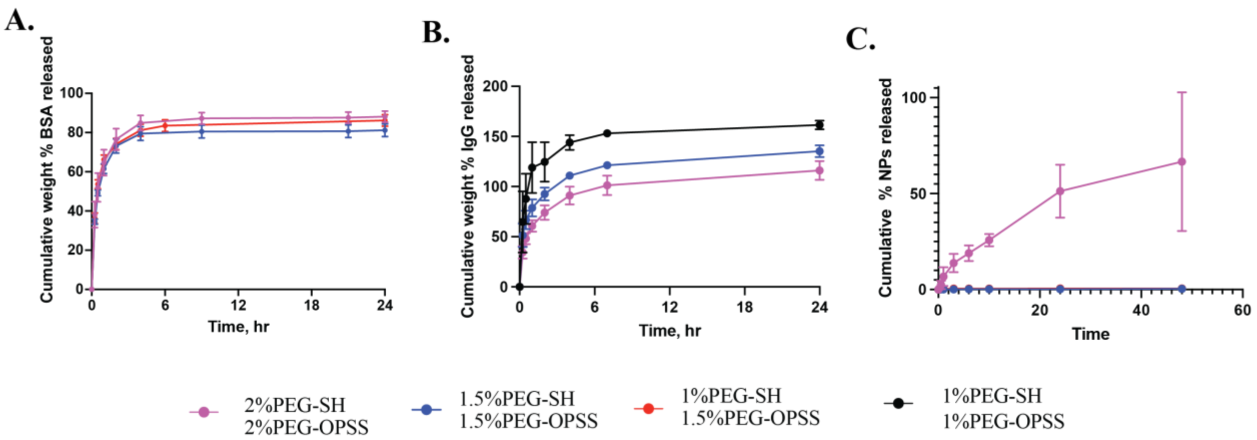


Figure S5. Release kinetics of model therapeutic cargo from different formulations of rapid forming PEG hydrogels. Cumulative release profile of (A) BSA from 10kD PEG gels (n=3), (B) IgG from 10kD PEG gels (n=3), and (C) 20nm nanoparticles (NP) from 20kD PEG gels (n=3) over 24 hours.

FITC labeled IgG was used for these release experiments. More than 100% release was observed likely due to fluorescence dequenching following IgG hydrolysis over time.[1]

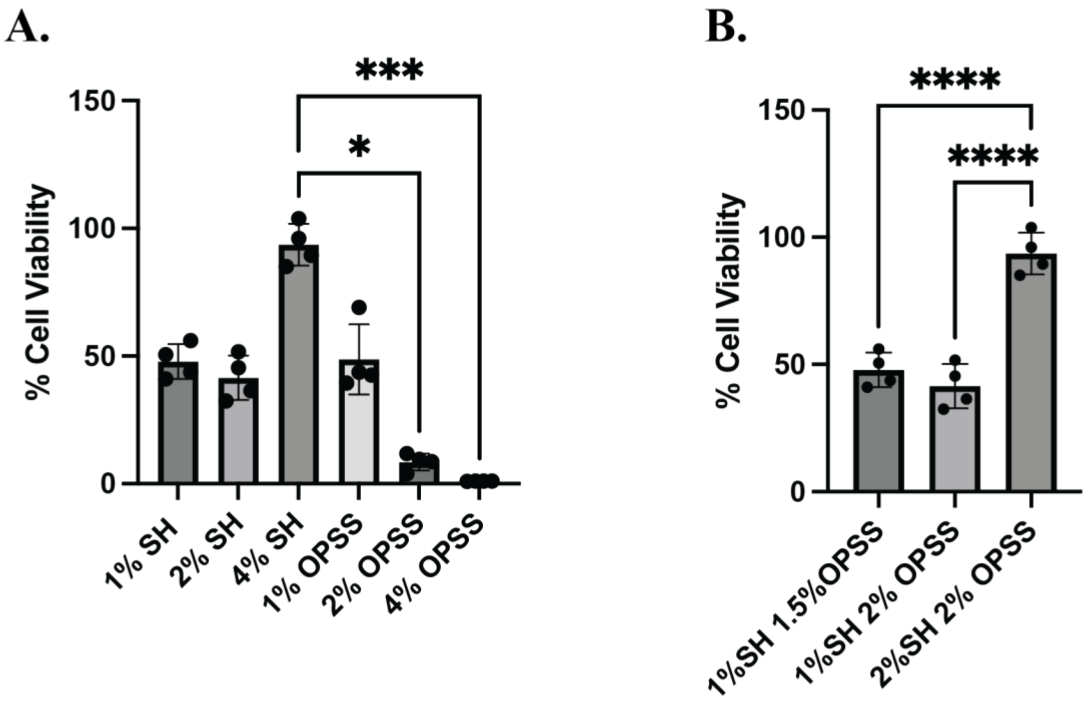


Figure S6. Biocompatibility of rapid forming 5kD PEG gels. Viability of HEK-293T cells following treatment with (A) 4-arm PEG solutions (n=4) * $p$ <0.05 ***$p$ <0.001 for Kruskal-Wallis test. (B) PEG hydrogels (n=4) ****$p$ <0.0001 for one-way ANOVA with Tukey's multiple comparison test.

References:

[1] C. Wischke, H.-H. Borchert, Fluorescein isothiocyanate labelled bovine serum albumin (FITC-BSA) as a model protein drug: opportunities and drawbacks, Pharmazie 61 (2006) 770–774.
